# Transposable element accumulation drives genome size increase in *Hylesia metabus* (Lepidoptera: Saturniidae), an urticating moth species from South America

**DOI:** 10.1101/2024.07.11.602864

**Authors:** Charles Perrier, Rémi Allio, Fabrice Legeai, Mathieu Gautier, Frédéric Bénéluz, William Marande, Anthony Theron, Nathalie Rodde, Melfran Herrera, Laure Saune, Hugues Parrinello, Melanie Mcclure, Mónica Arias

## Abstract

We present the first nuclear genome assembly and a complete mitogenome for *Hylesia metabus* (Arthropoda; Insecta; Lepidoptera; Saturniidae). The assembled nuclear genome sequence is 1,271 Mb long, which is among the 10 largest lepidopteran genome assemblies published to date. It is scaffolded in 31 pseudo chromosomes, has a BUSCO score of 99.5%, and has a highly conserved synteny compared to phylogenetically close species. Repetitive elements make up 67% of the nuclear genome and are mainly located in intergenic regions, among which LINEs were predominant, with CR1-Zenon being the most abundant. Phylogenetic and comparative analyses of *H. metabus* assembly and 17 additional Saturniidae and Sphingidae assemblies suggested that an accumulation of repetitive elements likely led to the increased size of *H. metabus’* genome. Gene annotation using Helixer identified 26,122 transcripts. The Z scaffold was identified using both a synteny analysis and variations of coverage for two resequenced male and female *H. metabus*. The *H. metabus* nuclear genome and mitogenome assemblies can be found and browsed on the BIPAA website and constitute useful resources for future population and comparative genomics studies.

## Introduction

The yellowtail moth *Hylesia metabus* (Saturniidae, Lepidoptera, Figure 1) known as “palometa peluda” in Venezuela and “papillon cendre” in French Guiana, is probably the most studied *Hylesia* species due to the health problems it causes. Like other species in the genus, adult females have urticating hairs that are easily released into the air, which can then come into contact with humans and cause a painful dermatitis (referred to as “Caripito itch” in Spanish or “papillonite” in French) and in extreme cases can cause respiratory problems (Rodriguez-Morales et al., 2005). Unlike other species, *H. metabus* is largely distributed in northern South America and it is responsible for epidemic outbreaks in Venezuela and French Guiana (Ciminera et al., 2019; Hernández et al., 2012; Jourdain et al., 2012). During outbreaks, hundreds to thousands of females fly simultaneously over human settlements, attracted by urban lights (Jourdain et al., 2012). The resulting abundance of urticating hairs negatively impact society by forcing citizens to shut themselves inside their houses at dusk so as to limit risks of dermatitis, and schools are forced to close to prevent children getting into contact with urticating hairs that remain on school grounds (ANSES French Agency for food environmental and occupational health & safety, 2011). *Hylesia metabus* populations are present in heterogeneous environments such as forest, savannahs and mangroves, although only populations in coastal areas are known to cause problems of epidemic dermatitis (Jourdain et al., 2012; Rodriguez-Morales et al., 2005). Using mitochondrial markers and nuclear microsatellite markers, previous studies have shown that, although *H. metabus* populations do belong to a single species, populations are genetically differentiated at relatively small spatial scale in French Guiana and Venezuela, notably between forest and mangrove habitats (Cequena et al., 2012; Ciminera et al., 2019). To better investigate the genomics of *H. metabus* populations and the potential genetic determinants responsible for a population’s propensity to produce problematic outbreaks, more in-depth genomics studies are needed. Hence, sequencing, assembling and annotating the first reference genome for *H. metabus* was essential to enable future population genomic studies, and here we achieved this using PacBio HiFi long reads scaffolded with Omni-C data.

**Figure 1.**
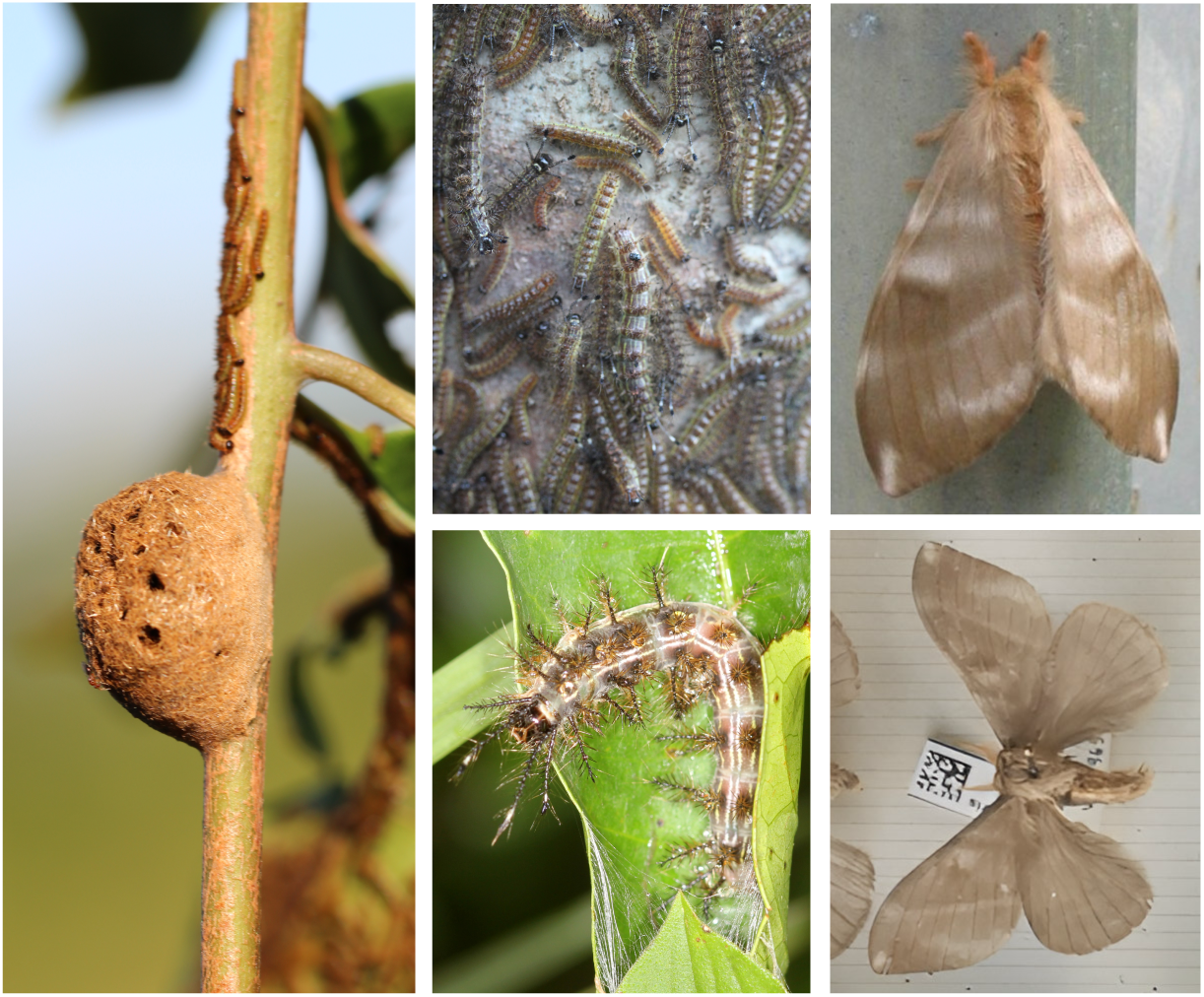
Pictures of *Hylesia metabus* at different developmental stages: nest covered by urticating hairs and with first stage larvae (left); gregarious larvae at stages L2, L3, L4 (upper middle); larva stage L7 (lower middle - stage used for DNA extraction); adult female (up right); adult male (lower right). (Photos of nest and larvae ©Jean-Philippe Champenois).

## Materials and methods

### Sample collection

In Stoupan (4.750 N 52.331 W), French Guiana, we collected in September 2021 two *H. metabus* larvae for the genome sequencing and scaffolding, with one larva being used for the HiFi library construction and the other larva used for the Omni-C library construction. We also collected 1 adult male and 1 adult female, for whole genome individual resequencing. Each larva was flash frozen in liquid nitrogen before being stored at -80°C. Each adult was stored in ethanol 85° and stored at -20°C.

### DNA extractions, libraries preparations and sequencing

For HiFi sequencing, high molecular weight (HMW) DNA was extracted from 0.3g of H. *metabus*, from the first larvae, using QIAGEN Genomic-tips 500/G kit (Qiagen, MD, USA). We followed the tissue protocol extraction, which in brief consisted of 0.3g of frozen *H. metabus* larvae abdomen ground in liquid nitrogen with a mortar and pestle. After 3h of lysis and one centrifugation step, the DNA was immobilized on the column. After several washing steps, DNA was eluted from the column, then desalted and concentrated by Isopropyl alcohol precipitation. A final wash in 70% ethanol was performed before resuspending the DNA in EB buffer. Analyses of DNA quantity and quality were performed using NanoDrop and Qubit (Thermo Fisher Scientific, MA, USA). DNA integrity was also assessed using the Agilent FP-1002 Genomic DNA 165 kb on the Femto Pulse system (Agilent, CA, USA). Hifi library was constructed using SMRTbell® Template Prep kit 2.0 (Pacific Biosciences, Menlo Park, CA, USA) according to PacBio recommendations (SMRTbell® express template prep kit 2.0 - PN: 100-938-900). HMW DNA samples were first purified with 1X Agencourt AMPure XP beads (Beckman Coulter, Inc, CA USA), and sheared with Megaruptor 3 (Diagenode, Liège, BELGIUM) at an average size of 20 kb. After End repair, A-tailing and ligation of SMRTbell adapter, the library was selected on BluePippin System (Sage Science, MA,USA) for a range size of 10-50kb. The size and concentration of the library were assessed using the Agilent FP-1002 Genomic DNA 165 kb on the Femto Pulse system and the Qubit dsDNA HS reagents Assay kit. Sequencing primer v5 and Sequel® II DNA Polymerase 2.2 were annealed and bound, respectively, to the SMRTbell library. The library was loaded on one SMRTcell 8M at an on-plate concentration of 90pM. Sequencing was performed on the Sequel® II system at Gentyane Genomic Platform (INRAE Clermont-Ferrand, France) with Sequel® II Sequencing kit 3.0, a run movie time of 30 hours with an Adaptive Loading target (P1 + P2) at 0.75. After filtering and correcting the SMRTcell output, we obtained 2,117,541 reads totalizing 42.7 Gb, with an N50 of 20,851 bp measured with LongQC v1.2.1 (Fukasawa et al., 2020) .

The Omni-C library (Dovetail Genomics®) was produced according to the manufacturer instructions. In brief, this consisted of 33mg of frozen *H. metabus* abdomen of the second larva, ground in liquid nitrogen and suspended in PBS. Then the DNA was fixed with formaldehyde and digested using 2µl of a nuclease enzyme mix. After binding 500ng of the digested DNA to chromatin capture beads, a proximity ligation was performed and the crosslinks were reversed to produce the linked DNA. Finally, 107ng of the linked DNA was used to produce a library then paired-end (2×150bp) sequenced on three different S4 flow-cell lanes on an Illumina® NovaSeq system. The obtained raw paired-end reads were filtered using fastp v0.23.2 (Chen et al., 2018) run with default options leading to a total of 171 millions of read pairs (51.2 Gb).

Two whole genome libraries of one adult female and one adult male were constructed using truseq kit to produce illumina short reads. In brief, DNA from one adult female and one adult male was extracted using the blood and tissue kit, column style, from Qiagen (Qiagen, MD, USA). DNA concentration was estimated using both Qubit fluorometric measures and nanodrop absorbance measures. Libraries were then constructed using the TruSeq nano DNA kit from Illumina and paired-end (2×150bp) sequenced on a S4 flow-cell lanes on an Illumina® NovaSeq system, producing 234 millions read pairs for the male and 325 millions read pairs for the female after filtering with fastp v0.23.2.

### Nuclear genome assembly, filtering and scaffolding

We used Jellyfish (Marçais & Kingsford, 2011) and GenomeScope (Ranallo-Benavidez et al., 2020; Vurture et al., 2017) to estimate the genome size from HiFi data. A whole genome assembly was then built from HiFi reads HiFiasm v0.16.1 (Cheng et al., 2021) ran with default options.

We detected and filtered out the mitogenome, potential contaminants and haplotigs as follows. Using MitoFinder v1.4.2 (Allio et al., 2020) on the primary assembly, we detected and annotated a contig corresponding to the complete mitochondrial genome sequence of *H. metabus*. The mitochondrial genome was removed from the nuclear genome assembly and we drew a circular representation using CGView (Stothard & Wishart, 2005). We used blobtools (Laetsch & Blaxter, 2017) to filter out potential contaminants. We also searched for potential contaminant sequences originating from the plant species’ genome on which larvae were found feeding on (*Tapirira guianensis*) by mapping short reads from this species (https://www.ebi.ac.uk/ena/browser/view/ERR7620141) on our putative *H. metabus* scaffolds using bwa-mem2 v2.2.1 (Vasimuddin et al., 2019) and samtools v1.10 (Danecek et al., 2021). We used purge haplotigs v1.1.2 (Roach et al., 2018) to remove potential haplotigs and small contigs exhibiting bad mapping quality or that could be considered as junk or repeats.

The contig assembly was scaffolded using Omni-C sequencing data. Following Serizay et al. (2024), we separately mapped the filtered paired-end reads from the three sequencing lanes to the contig assembly using bwa-mem2 v2.2.1 (Vasimuddin et al., 2019) which was run with options *-SP5M*. The three resulting bam files generated with the samtools v. 1.14 *view* (Li et al., 2009) were then parsed, sorted and deduplicated with *pairtools v1.0.3* (Open2C et al., 2023) programs *parse* (run with options –*min-mapq 20* and *—drop-sam*), *sort,* and *dedup*, respectively. The three *pairs* files were further merged with the *pairtools merge* program and the *dump* program from the *cooler* v0.9.3 suite (Abdennur & Mirny, 2020) was used to generate a contact matrix (using 500 kb bins) that was visualized using a custom R function. The identified Omni-C pairs were finally used to scaffold the assembly using *YaHS v1.2* (Zhou et al., 2023) ran with default options but *–file-type PA5* specification to read the *pairtools* generated pair file. A contact map for the resulting scaffolded assembly was generated as described above after converting Omni-C pairs mapping coordinates from the contig assembly (based on the agp file) using a custom *awk* script.

Completeness of the contig and scaffolded assemblies was evaluated using Benchmark Universal Single Copy Orthologs (BUSCO v5.5.0, Manni et al., 2021; Simão et al., 2015) for “arthropoda_odb10” and “lepidoptera_odb10” databases. Blobtools (Laetsch & Blaxter, 2017) was used to draw a snailplot graph.

### Annotation of protein coding genes and repetitive elements

Protein coding gene prediction was achieved on the non-masked genome version of *H. metabus*. Helixer v0.3.0 with the option --lineage invertebrate was used for gene prediction (Holst et al., 2023). The quality of the annotation was assessed with BUSCO v5.2.2 using lineage dataset lipidoptera_odb10, PSAURON 1.0.2 (Sommer et al., 2024) and OMArk 2023.10 (Nevers et al., 2024). Functional annotation of the protein sequences obtained with GFFread (Pertea & Pertea, 2020) from the Helixer output were done with Diamond v2.0.13 (Buchfink et al., 2015) on NCBI NR 2022-12-11, Blast2GO Command Line v1.5.1 (Götz et al., 2008), eggNOG v2.1.9 (Huerta-Cepas et al., 2019) with eggnog database v5.0.2 and Interproscan v5.59-91.0 (Jones et al., 2014). The genome sequence and its annotations can be browsed at https://bipaa.genouest.org/sp/hylesia_metabus/.

Repetitive elements were identified using EarlGrey (Baril et al., 2024) v4.1.0, which notably uses RepeatMasker (Smit et al., 2015), RepeatModeler2 (Flynn et al., 2020), and LTR_Finder (Xu & Wang, 2007). We used the Arthropoda repeat library from DFAM 3.5 as the initial repeats library. We inspected the distribution of repetitive elements across the genome, in intergenic regions, introns, UTR, and CDS from the Helixer GFF file.

### Phylogenetic and comparative analyses

Phylogenetic analyses have been done by comparing the BUSCO sequences from the genome of *H. metabus* to the one of 4 other Saturniidae species (*Automeris io, Samia ricini, Antheraea yamamai, Saturnia pavonia*), 12 Sphingidae species (*Cephonodes hylas, Hemaris fuciformis, Hyles euphorbiae, Deilephila porcellus, Theretra japonica, Manduca sexta, Lapara coniferarum, Sphinx pinastri, Clanis bilineata, Laothoe populi, Amorpha juglandis, Mimas tiliae*), and *Bombyx mori* as outgroup. External assemblies were downloaded from NCBI using their respective accession number with the command-line tool “datasets” (e.g. datasets download genome accession GCF_030269925.1 --filename Bombyx_mori.zip; see supplementary material 1 for accessions). For each assembly, BUSCO sequences were annotated and extracted based on the predefined dataset “Lepidoptera_odb10”, which includes 5,286 orthologous genes for Lepidoptera. BUSCO search was performed through the gVolante web server (Nishimura et al., 2017). Amino acid sequences corresponding to every BUSCO marker were first individually aligned with the MAFFT (Katoh & Standley, 2016) algorithm FFT-NS-2. All markers were then concatenated in one supermatrix using seqCat.pl and seqConverter.pl (Bininda-Emonds, 2006; Fasterius & Al-Khalili Szigyarto, 2019). Phylogenetic inferences were performed with Maximum-likelihood (ML) as implemented in IQ-TREE V2.2.2.6 (Minh et al., 2022). One partition per BUSCO sequence was defined with the option “-spp”. The best evolutionary model was selected for each partition using ModelFinder implemented in IQ-TREE and some partitions were merged if necessary (-m MFP+MERGE). Following the recommendation of IQ-TREE developers, we also set a smaller perturbation strength (-pers 0.2) and a larger number of stop iterations (-nstop 500) to avoid local optima. Finally, node supports were evaluated with UltraFast Bootstraps (UFBS) estimated by IQ-TREE (-bb 1000). UFBS are considered robust when higher than 95%.

Repetitive elements content and repeat landscape in *H. metabus* was compared to the one in the 17 species mentioned above, for which we ran the same EarlGrey pipeline as for *H. metabus*.

Synteny between the genome of *H. metabus* and the genomes of *Antheraea yamamai, Saturnia pavonia*, *Deilephila porcellus, Manduca sexta, Laothoe populi,* and *Bombyx mori, extracted from NCBI (see above and supplementary table 1)* were inspected with genespace (Lovell et al., 2022) using the gene positions derived from the GFF files, and their corresponding orthogroups predicted with orthofinder v2.5.5 (Emms & Kelly, 2019). Because protein coding genes annotation of *Antheraea yamamai, Saturnia pavonia*, *Deilephila porcellus,* and *Laothoe populi* were not available at NCBI, we produced their respective annotations with Helixer (v0.3.3, with the options --lineage invertebrate --subsequence-length 108000 --overlap-offset 54000 --overlap-core-length 81000). Finally, the sizes of intergenic and genic regions were calculated from the GFF file with a custom script.

### Identification of the Z scaffold

In order to identify the scaffold corresponding to the Z chromosome in *H. metabus*, we first inspected synteny graphs with *Deilephila porcellus* and *Saturnia pavonia* assemblies for which the Z scaffold was known. We also used Dgenies (Cabanettes & Klopp, 2018) to examine more precisely the synteny between *Hylesia metabus* and *Deilephila porcellus* (Boyes et al., 2022). Second, we investigated potential coverage variation among scaffolds between the resequenced male and female. To do so, we first used Fastp (Chen et al., 2018) to keep only good quality sequences that we then aligned to the *H. metabus* assembly using bwa-mem2 v2.2.1 (Vasimuddin et al., 2019). We then used samtools to sort and index reads and to estimate coverage per scaffold for each individual (Danecek et al., 2021). For each scaffold we measured the female to male ratio of percentage of reads mapped to each scaffold. A ratio of approximately 50% would indicate the Z scaffolds, while ratios of 100% would indicate autosomal scaffolds. Finally, in order to determine the genotypic sex of the individual larvae sequenced to assemble the genome, we investigated potential variations of HiFi raw reads coverage between the putative Z scaffold and the other large scaffolds. A coverage deficit of about 50% on the Z scaffold would illustrate that the sequenced individual was a female.

## Results and discussion

### Assembly of a 1.27 GB long genome scaffolded in 31 pseudo chromosomes

The Jellyfish and Genomescope analysis of the HiFi reads suggested that the genome size was 1.19 Gb with a heterozygosity of 2.19% (Figure 2A). The Hifiasm assembly consisted of 171 contigs ranging from 15kb to 57Mb and totalizing 1.41 Gb, with an N50 of 34,63 Mb, an L50 of 17 and 39.85% of GC (Supplementary material 2). MitoFinder identified the smallest (15393 bp) and most covered (174 X) contig, as being the complete mitochondrial genome sequence and annotated it fully (Supplementary material 3). The final assembly, after decontamination, purge of haplotigs, removal of the mitochondrial contig, and scaffolding with Omni-C, consisted in 31 scaffolds totalizing 1.27 Gb, with an N50 of 45,18 Mb, an L50 of 14 and 39.58% of GC (Figure 2B, Supplementary material 4). The contact map of the primary assembly showed very high contiguity even before scaffolding (Supplementary material 5), enabling an efficient scaffolding toward a chromosome level (Figure 2C). The haploid read depth was on average 14 X, slightly lower than the targeted read depth, as a consequence of a larger genome size than expected. Arthropoda BUSCO gene representation of the final assembly was 99.5% complete with 98.7% single-copy genes, and the Lepidoptera BUSCO gene representation was 98.6% complete with 97.6% single-copy genes (Supplementary material 4, Figure 2B), with less than 1% of duplicated genes.

**Figure 2.**
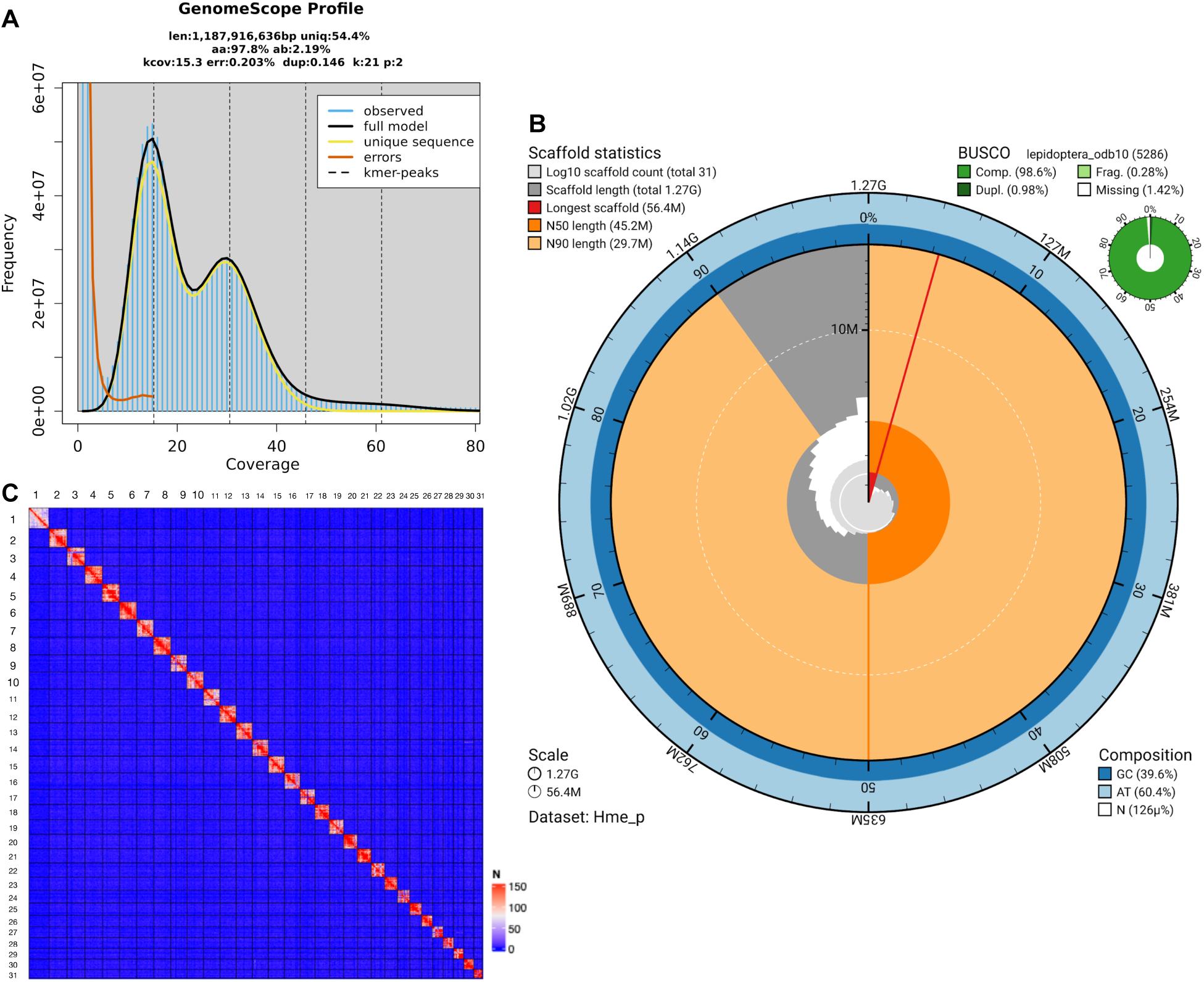
A) K-mer spectra output generated from corrected PacBio HiFi data using GenomeScope. The bimodal pattern observed corresponds to a diploid heterozygous genome. B) BlobToolKit Snailplot showing N50 metrics and BUSCO gene completeness. C) Omni-C contact map of the scaffolded genome sequence of *Hylesia metabus* (number of contacts per 500 kb bin).

### Highly conserved synteny but size increase of intergenic regions contributing to a large genome size compared to phylogenetically close species

Synteny was highly conserved between *H. metabus* and phylogenetically close species (Figure 3A), suggesting no evidence for large chromosomal rearrangements and good quality of the assembly. In addition, the number of scaffolds for our assembly was identical to *Saturnia pavonia*, and very similar to other close species, indicating no chromosomal fusion or fission.

**Figure 3.**
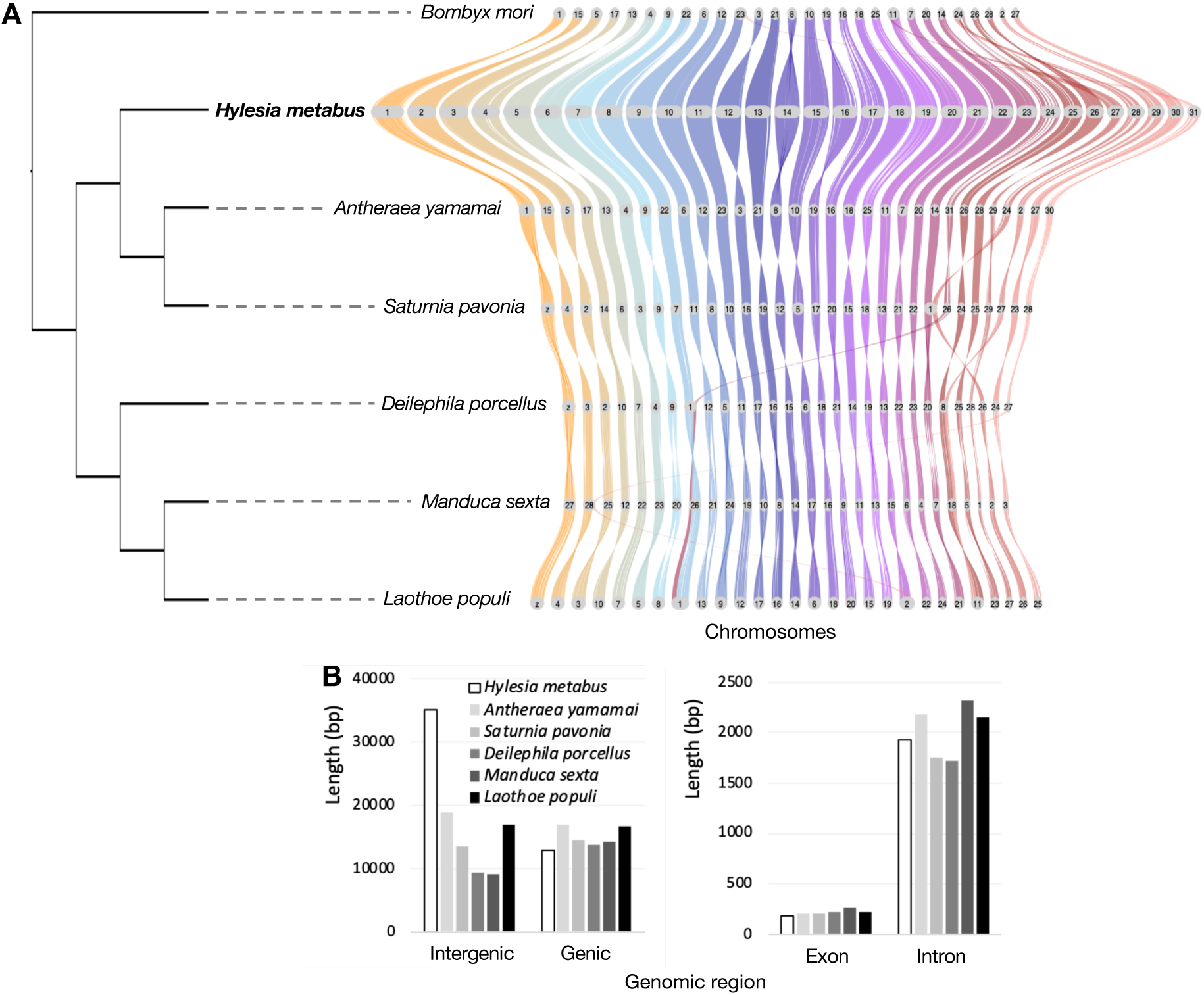
A) Phylogeny and synteny for *Hylesia metabus* and 2 other Saturniidae species, 3 Sphingidae species, and *Bombyx mori*. B) Length of intergenic and genic regions, and of exons and introns, for *H. metabus* and 2 other Saturniidae species, 3 Sphingidae species, and *Bombyx mori*.

However, the genome sequence size, 1.27 Gb, was much larger than for other available Saturniidae genome assemblies. For example, the genome assemblies for the close species *Automeris io* and *Saturnia pavonia* were both 490 Mb long (Crowley et al., 2024; Skojec et al., 2024). Yet, other lepidoptera species are known to have large genomes, notably *Euclidia mi*, 2.32 GB (Boyes & Holland, 2023), *Parnassius behrii* 1.59 GB (GCA_036936625.1), *Parnassius apollo* 1.4 GB (Podsiadlowski et al., 2021), *Tholera decimalis* 1.33 GB (Boyes et al., 2023), *Thaumatotibia leucotreta*, 1.28 GB (Bierman et al., 2023), *Graphium colonna* 1.27 GB (Triant & Pirro, 2023). To date, the *H. metabus* genome is amongst the top 10 largest lepidopteran genome assemblies present on NCBI database as of 24/05/2024. In line with the highly conserved synteny, the increase in total genome size, and the very low level of duplicated BUSCO genes, we found much larger intergenic regions in *H. metabus* compared to phylogenetically close species but comparable sizes of genic regions (Figure 3B), both for introns and exons, and in a regular manner across chromosomes (Supplementary material 6).

### Invasion of repetitive elements

Analysis of repetitive elements was achieved with the fully automated EarlGrey pipeline and determined that 67% of the *H. metabus* genome sequence was made-up of repeats (Figure 4A & 4B, Supplementary material 1 & 4). Compared to other species of Saturniidae and species of Sphingidae, *H. metabus* had a higher proportion of repetitive elements (Figure 4A, Supplementary material 1 & 4), although this was similar to the 65% reported for the aforementioned large genome of *Parnassius apollo* (Podsiadlowski et al., 2021). In general, TE invasion in lepidoptera, and more broadly in arthropods, is associated with increased genome size (Gilbert et al., 2021; Muller et al., 2021; Petersen et al., 2019).

**Figure 4.**
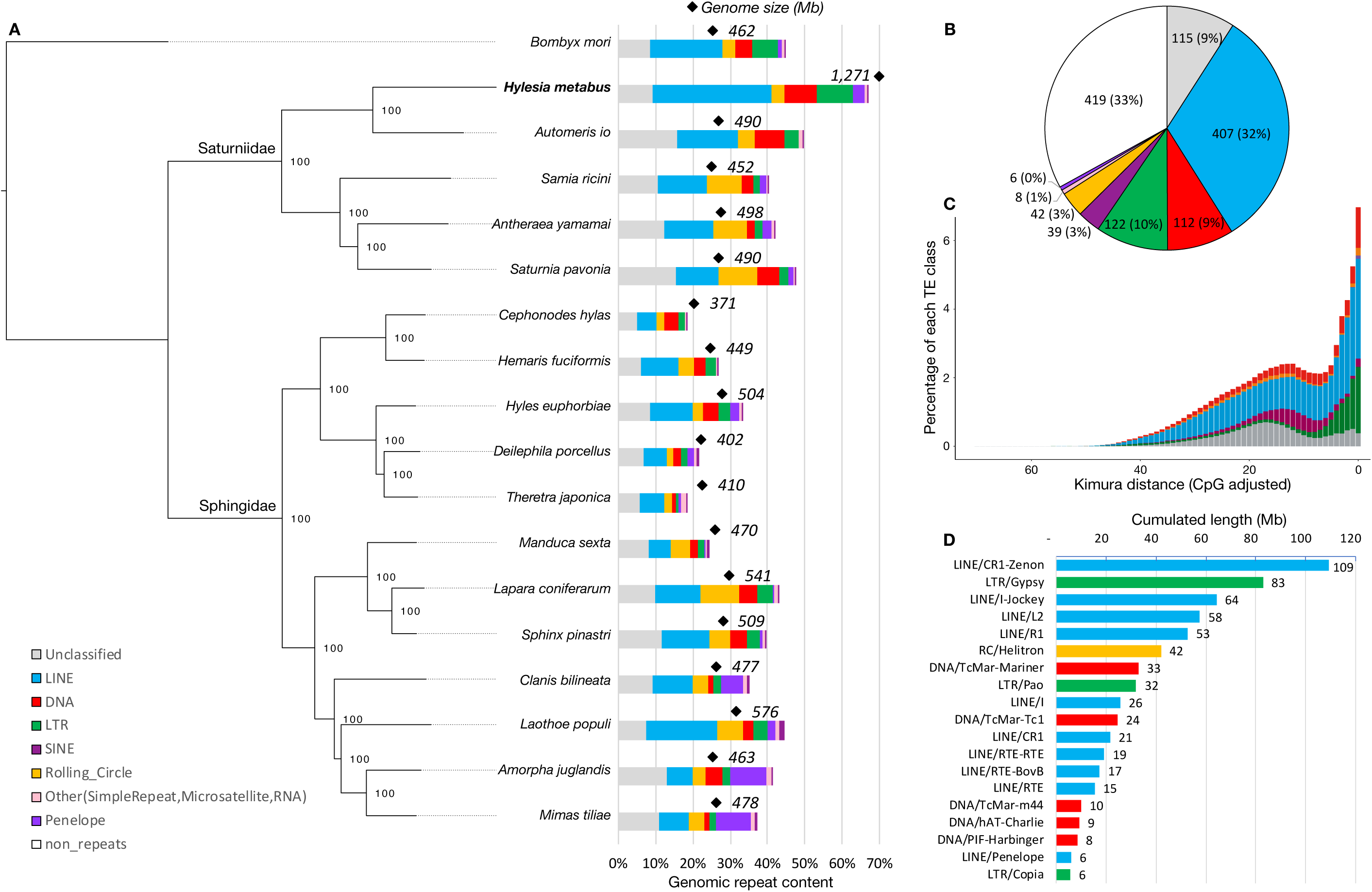
A) Phylogeny and comparative analysis of repeat content for *Hylesia metabus* and 4 other Saturniidae species, 12 Sphingidae species, and *Bombyx mori* as an outgroup. Phylogenetic tree (left panel), genome sequence length and repeat content (right panel). Repeat content (B), repeat landscape (C) and cumulated length for each of the most abundant TE families (D) in the genome assembly of *H. metabus*.

Repetitive elements were found more often in intergenic regions (77.1%) than in introns (21.5%), UTR (0.3%) and CDS (1.1%) (Supplementary material 7A). Moreover, intergenic regions consisted of repeats (68.1%) more than introns (60.4%), UTR (25.8%) and CDS (28.0%; Supplementary material 7B). This reinforces the hypothesis that the invasion of repetitive elements explains the large size of intergenic regions and of the entire genome in *H. metabus*.

The majority of repetitive elements identified in *H. metabus* were of type LINE (32% of the genome, hence 55% of the classified repeats; Figure 4B), followed by LTR, DNA and SINE elements (respectively 10%, 9% and 3% of the genome, hence 17%, 16% and 5% of the classified repeats). Only 9% of the genome was made of unclassified repetitive elements. These LINEs and LTRs proportions are higher than for other species of Saturniidae and species of Sphingidae (Figure 4A, Supplementary material 1). This higher proportion of LINEs and LTRs is comparable to *Parnassius apollo*.

The repeat landscape of *H. metabus* suggests that repetitive elements, especially of type LINE, invaded the genome relatively regularly over time, but that LTRs had a recent burst of invasion (Figure 4C). While comparing the repeat landscape of *H. metabus* to the 4 other species of Saturniidae and 12 species of Sphingidae (Supplementary material 8), we found that repeat landscapes were in general relatively similar between close species and dissimilar between more distant ones. For example, while the repeat landscape of *H. metabus* was very similar to the one of its closest species, *Automeris io*, it was very dissimilar to the repeats landscapes of the three other Saturniidae species *Samia ricini*, *Antheraea yamamai* and *Saturnia pavonia* which all showed a more ancient burst of invasion and a higher accumulation of Rolling circles. Regarding the timing of accumulation of these transposable elements, under uncorrelated relaxed clock model, Rougerie et al 2022 estimated the origin of the crown group *Hylesia* at 10 to 13 MY (unfortunately *H. metabus* was not included in their study), the divergence time between the genera *Hylesia* and *Automeris* at about 25 MY and about 46 MY between *Hylesia* and the genera *Samia*, *Antheraea* and *Saturnia*. Skojec et al. (2024) study found similar divergence times (although showing a 18MY divergence between *Automeris* and *io* clades). This suggests that the similar repeat landscapes of *H. metabus* and *Automeris io* might have evolved between 46 and 25 MY, and that *H. metabus* accumulation of TE might have occurred during the last 25 MY, and perhaps more likely during the *Hylesia* genus diversification between 10 and 13 MY or during the more recent evolution of *H. metabus*. Sequencing genomes for more Saturniidae species, especially from the genus *Hylesia*, would enable more detailed comparative analyses of the accumulation of transposable elements in this family.

The five most abundant repetitive elements were LINE/CR1-Zenon (109M bp - 319,358 copies, Figure 4D, Supplementary material 9), LTR/Gypsy (83M bp), LINE/I-Jockey (64M bp), LINE/L2 (58M bp) and LINE/R1 (42M bp). For the four other Saturniidae species considered in this study (Supplementary material 10), LINE/R1 was the most abundant family in *Automeris io,* followed by RC/Helitron and LINE/CR1-Zenon, and RC/Helitron was the most abundant family in the three other species, followed by LINE/I-Jockey in two species and by LINE/L2 and LINE/R1 in the last one. This more detailed analysis therefore shows that the TE invasion of *H. metabus* genome is primarily driven by a few TE families that are also among the most abundant ones in the closest species genome sequences. In particular, the LINE CR1 *Zenon* has been shown to successfully invade other lepidopteran species’ genomes (Wang et al., 2019).

### Gene prediction, Z scaffold and data accessibility

Gene prediction with Helixer identified 26,122 transcripts on the non-masked version of the genome (Supplementary material 4). The BUSCO score for the annotated genes was 88.9% complete (Supplementary material 4). Comparatively OMArk identified 6361 complete proteins among the 6779 Obtectomera HOG, including 1466 duplicated, in the range of other closely related species. OMArk also determined that 77.25% of the protein sequences were placed at a consistent lineage, while 16.31% were unknown and reported no contamination. The psauron score, reflecting the likelihood of being a genuine protein coding sequence, was 96.9. Generating RNAseq data in order to further annotate this genome would probably greatly improve the precision of the annotation, by keeping only transcripts with evidence of expression.

The synteny inspection showed that the largest *H. metabus* scaffold (scaffold_1) corresponded to the Z scaffold in *Deilephila porcellus* and *Saturnia pavonia* assemblies (Figure 3A, Supplementary material 11). This was confirmed by the comparisons of read mapping between one male and one female (that had average coverage of 16X and 25X, respectively), revealing a 0.54 female to male relative ratio of coverage on scaffold 1 (Supplementary material 12). Finally, mapping HiFi reads of the assembled individual on the final assembly, we estimated that Scaffold_1 had on average 57% lower read depth than the other 30 largest scaffolds, furthermore confirming that scaffold_1 indeed corresponds to the Z chromosome and suggesting that the individual used to assemble the genome was a female.

## Conclusion

Here we present a high-quality genome sequence assembly for *H. metabus*, a Saturniidae moth species known for causing painful human dermatitis especially during recurrent demographic outbreaks. This genome sequence is among the 10 largest lepidopteran genome sequences published to date. The genome expansion could be explained by an invasion of repetitive elements, especially of LINEs and LTRs, in intergenic regions, as observed in a few other lepidopteran species. Both genome size, intergenic regions length and repeat content contrast with the closest species of Saturniidae and Sphingidae sequenced so far. It will be interesting to use several of the numerous *Hylesia* species (more than 110 species (Lemaire, 2002)) and additional Saturniidae species (Hamilton et al., 2019; Rougerie et al., 2022) as models to study repetitive element dynamics, notably by comparing their repeat contents, genomes sizes, genetic diversity and effective population sizes. Studying other *Hylesia* species genomes is also important as several of these species also result in health problems (Glasser et al., 1993; Iserhard et al., 2007; Molina, 2019; Salomón et al., 2005) and/or agricultural damage (Carrillo-Sánchez et al., 1998; Fronza et al., 2011). The genomic resource presented here will also be useful for future comparative studies of urticating insects’ genomes (Battisti et al., 2011), and for future population genomic studies of *H. metabus* aiming to better understand differences in population genetics and demography of this species in South America (Ciminera et al., 2019).

## Data availability

The nuclear and mitochondrial assemblies, and the HiFi, Omni-C and resequencing data will soon be available on NCBI under the BioProject ID PRJNA1132489. The nuclear and mitochondrial assemblies will also soon be available on the Bioinformatics BIPAA Platform, together with annotations of genes and repetitive elements, at the following address: https://bipaa.genouest.org/sp/hylesia_metabus/.

## Authors contributions

Conceptualization: C. Perrier, M. Arias; Funding acquisition: C. Perrier, M. McClure, M. Arias; Biological samples: F. Bénéluz; Pictures: M. Herrera; Wet lab and sequencing: C. Perrier, W. Marrande, A. Theron, N. Rodde, L. Sauné, H. Parrinello, M. Arias; Statistical analysis and visualization: C. Perrier, R. Allio, F. Legeai, M. Gautier, W. Marrande, M. Arias; Writing of the original draft: C. Perrier; Review & editing: All the authors.

## Acknowledgment

1. C. Perrier acknowledges INRAE CBGP for funding Omni-C analyses. M. McClure and M. Arias acknowledge CNRS-MITI (Mission pour les Initiatives Transverses) for funding HiFi analyses. We thank Genobioinfo and GenOuest INRAE platforms for giving access to bioinformatic computing facilities, Gentyane INRAE Genomic Platform for Pacific Biosciences sequencing, Pierre Nouhaud for advice on transposable element annotation, ARS Guyane in Cayenne for discussions regarding health issues caused by *H. metabus*, and Jean-Philippe Champenois for sharing pictures.

## Supplementary material

**Supplementary material 1.** Repeat composition, genome size and GenBank reference, for *Hylesia metabus* and 17 other species of saturniidae and sphingidae.

| Species | GenBank | Genome_size | LINE | Unclassified | Rolling_Circle | DNA | LTR | SINE | Other(SimpleRepeat, Microsatellite, RNA) | Penelope | Non_repeats |
| --- | --- | --- | --- | --- | --- | --- | --- | --- | --- | --- | --- |
| *Bombyx_mori* | GCF_030269925.1 | 461,704,601 | 19% | 8% | 4% | 4% | 7% | 1% | 0% | 0% | 55% |
| *Hylesia metabus* | not yet | 1,270,616,270 | 32% | 9% | 3% | 9% | 10% | 3% | 1% | 0% | 33% |
| *Automeris_io* | GCA_036320925.1 | 490212539 | 16% | 16% | 5% | 8% | 4% | 0% | 1% | 0% | 51% |
| *Samia_ricini* | GCA_014132275.2 | 452483983 | 13% | 11% | 9% | 3% | 2% | 2% | 0% | 0% | 60% |
| *Antheraea_yamamai* | GCA_036509395.1 | 498398989 | 13% | 12% | 9% | 2% | 2% | 3% | 0% | 0% | 58% |
| *Saturnia_pavonia* | GCA_947532125.1 | 489898868 | 11% | 15% | 10% | 6% | 3% | 1% | 0% | 0% | 53% |
| *Cephonodes_hylas* | GCA_030295005.1 | 371,083,271 | 5% | 5% | 2% | 4% | 2% | 0% | 0% | 0% | 82% |
| *Hemaris_fuciformis* | GCA_907164795.1 | 448,853,392 | 10% | 6% | 4% | 3% | 3% | 0% | 0% | 0% | 73% |
| *Hyles_euphorbiae* | GCA_023078785.2 | 504323440 | 12% | 8% | 3% | 4% | 3% | 2% | 0% | 0% | 67% |
| *Deilephila_porcellus* | GCA_905220455.2 | 402,071,895 | 6% | 7% | 2% | 2% | 2% | 2% | 1% | 0% | 78% |
| *Theretra_japonica* | GCA_033459515.1 | 409,552,430 | 7% | 6% | 2% | 1% | 1% | 1% | 1% | 0% | 82% |
| *Manduca sexta* | GCF_014839805.1 | 470036997 | 6% | 8% | 5% | 2% | 2% | 0% | 0% | 1% | 76% |
| *Lapara_coniferarum* | GCA_949316025.1 | 541279285 | 12% | 10% | 10% | 5% | 4% | 1% | 1% | 0% | 57% |
| *Sphinx_pinastri* | GCA_947568825.1 | 509,238,608 | 13% | 12% | 6% | 5% | 3% | 1% | 1% | 0% | 61% |
| *Clanis_bilineata* | GCA_036417725.1 | 477,454,959 | 10% | 9% | 4% | 1% | 2% | 6% | 1% | 1% | 65% |
| *Laothoe_populi* | GCA_905220505.1 | 576402658 | 19% | 8% | 7% | 3% | 4% | 2% | 1% | 1% | 55% |
| *Amorpha_juglandis* | GCA_949126905.1 | 463,042,227 | 7% | 13% | 4% | 5% | 2% | 10% | 1% | 0% | 59% |
| *Mimas_tiliae* | GCA_905332985.1 | 477,981,037 | 8% | 11% | 4% | 1% | 2% | 9% | 1% | 1% | 63% |

**Supplementary material 2.**
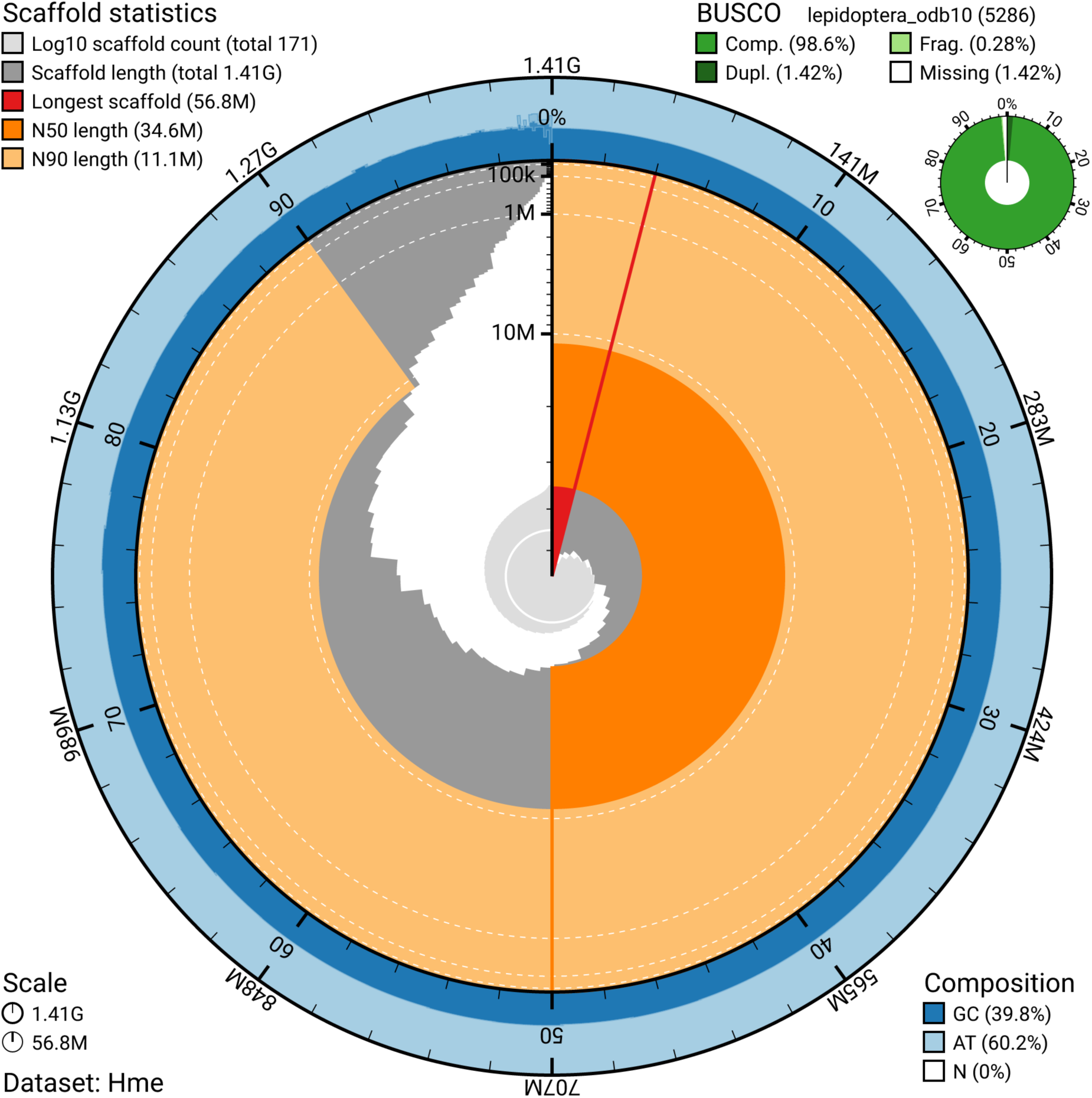
Snailplot on primary output hifiasm before scaffolding and decontamination.

**Supplementary material 3.**
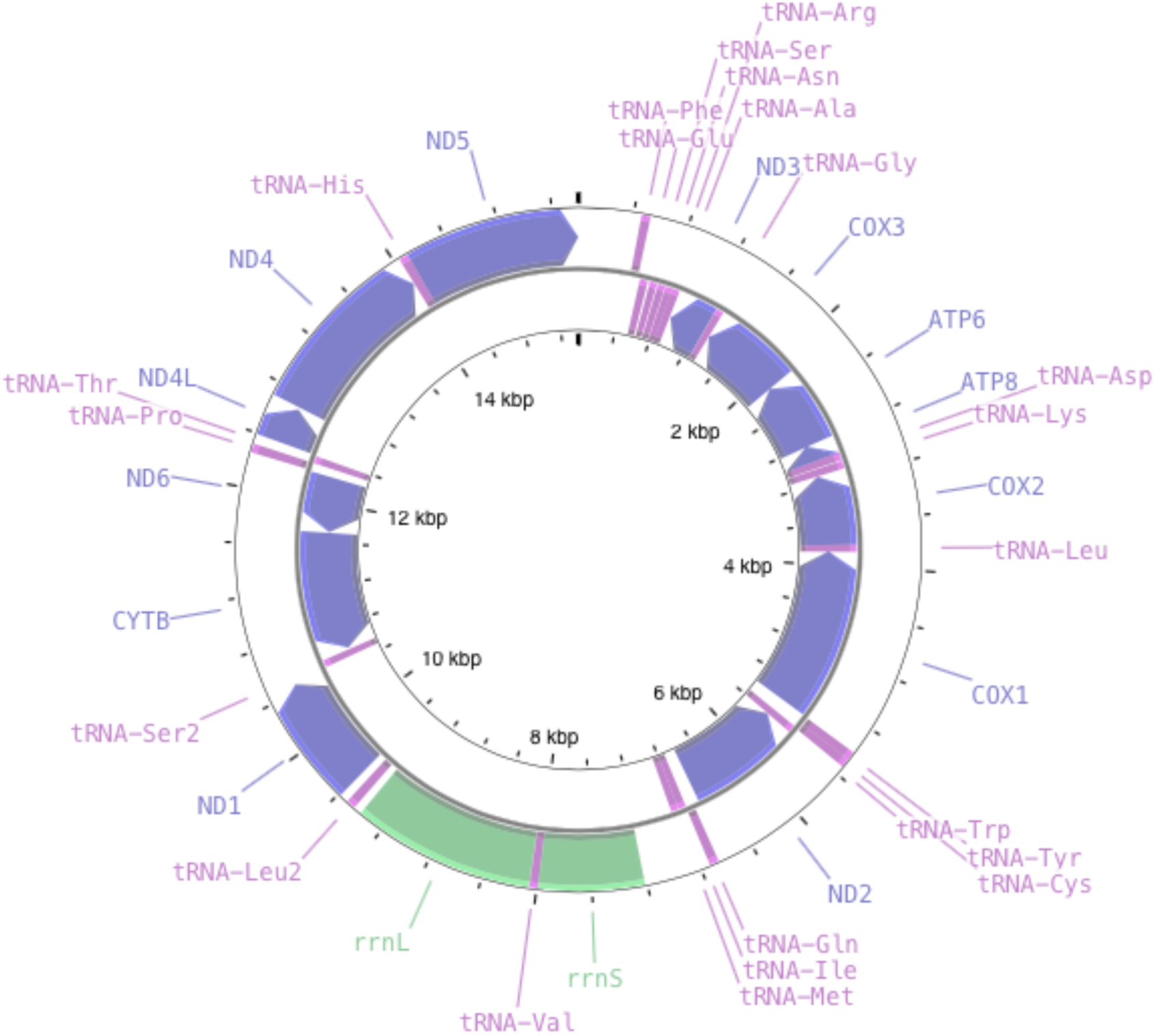
Mitogenome of *H. metabus*.

**Supplementary material 4.** Statistics of the assembly and annotations.

|  |  |  |  |
| --- | --- | --- | --- |
| Assembly statistics | n scaffolds |  | 31 |
|  | Largest scaffold |  | 56,404,000 |
|  | Total length of the scaffolded version |  | 1,270,616,270 bp |
|  | Estimated genome size |  | 1,187,916,636 bp |
|  | GC (%) |  | 39.58 |
|  | N50 for scaffolds |  | 45,175,336 |
|  | N90 for scaffolds |  | 29,682,000 |
|  | n N's per 100 kbp in the scaffolded version |  | 0.13 |
|  | BUSCO Arthropoda |  | C:99.5%[S:98.7%,D:0.8%],F:0.0%,M:0.5%,n:1013 |
|  | BUSCO Lepidoptera |  | C:98.6%[S:97.6%,D:1.0%],F:0.3%,M:1.1%,n:5286 |
| Annotation | Repeat content | all repeats | 67% |
|  |  | LINE | 32% |
|  |  | SINE | 3% |
|  |  | DNA | 9% |
|  |  | LTR | 10% |
|  |  | Unclassified | 9% |
|  | Gene prediction | n protein coding genes | 26122 |
|  |  | n transcripts | 26122 |
|  |  | gene size (mean / median bp) | 12942 / 11169 |
|  |  | transcript size (mean / median bp) | 1079 / 705 |
|  |  | CDS size (mean / median bp) | 847 / 517 |
|  |  | exon size (mean / median bp ) | 184 / 126 |
|  |  | intron size (mean / median bp) | 1934 / 966 |
|  |  | n exon per gene (mean / median) | 6.85 / 5 |
|  |  | BUSCO lepidoptera | C:88.9%[S:86.6%,D:2.3%],F:5.1%,M:6.0%,n:5286 |
|  |  | OMArk Completeness | 93.83% (S = 72.21%, D = 21.63%, M = 6.17%) |
|  |  | OMArk proteome assessment | Consistent lineage placement = 77.25%<br>Inconsistent lineage placement = 6.44%<br>Contamination = 0.0%<br>Unknown = 16.31% |

**Supplementary material 5.**
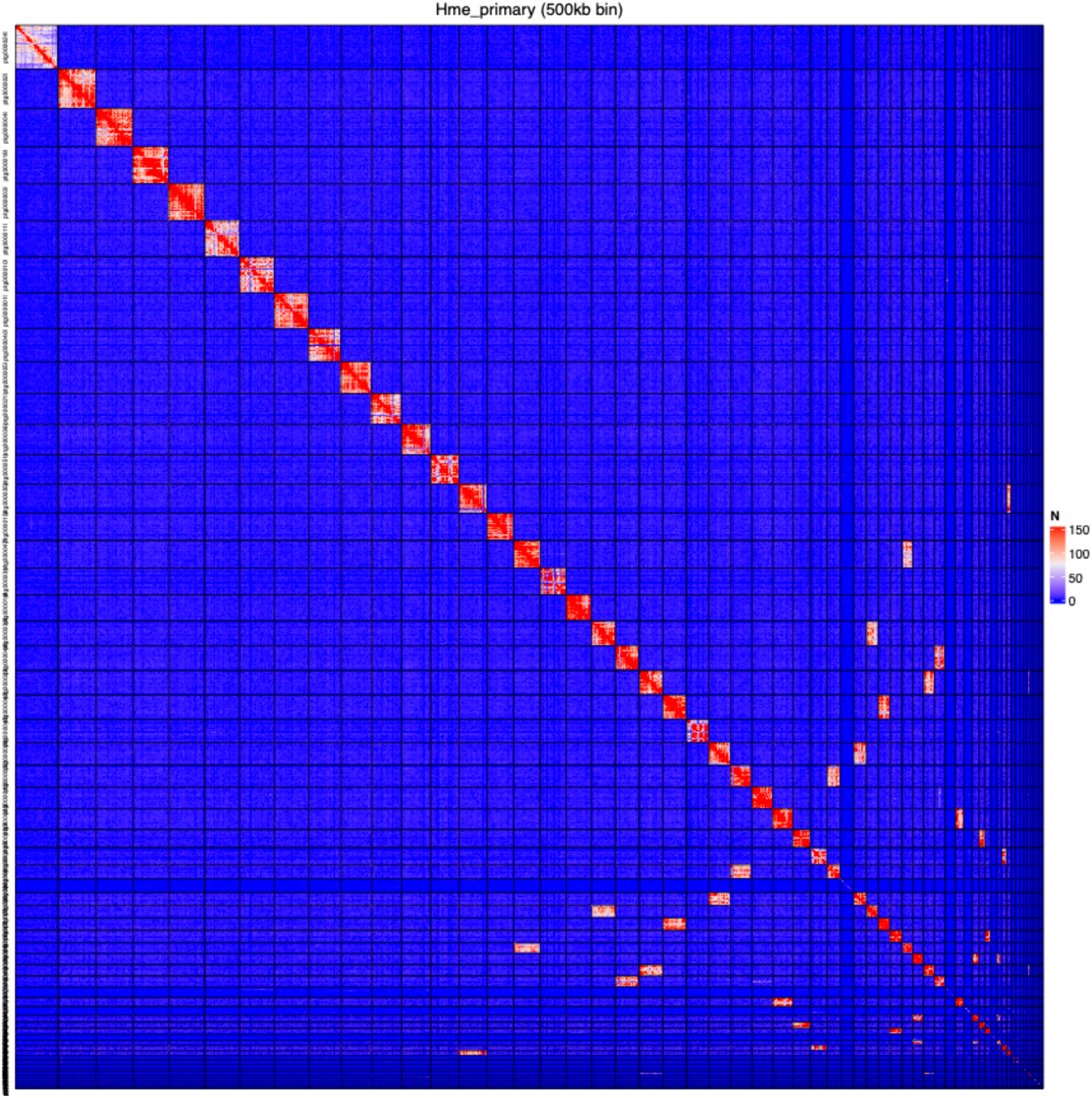
Contact map on primary contigs before scaffolding and decontamination.

**Supplementary material 6.**
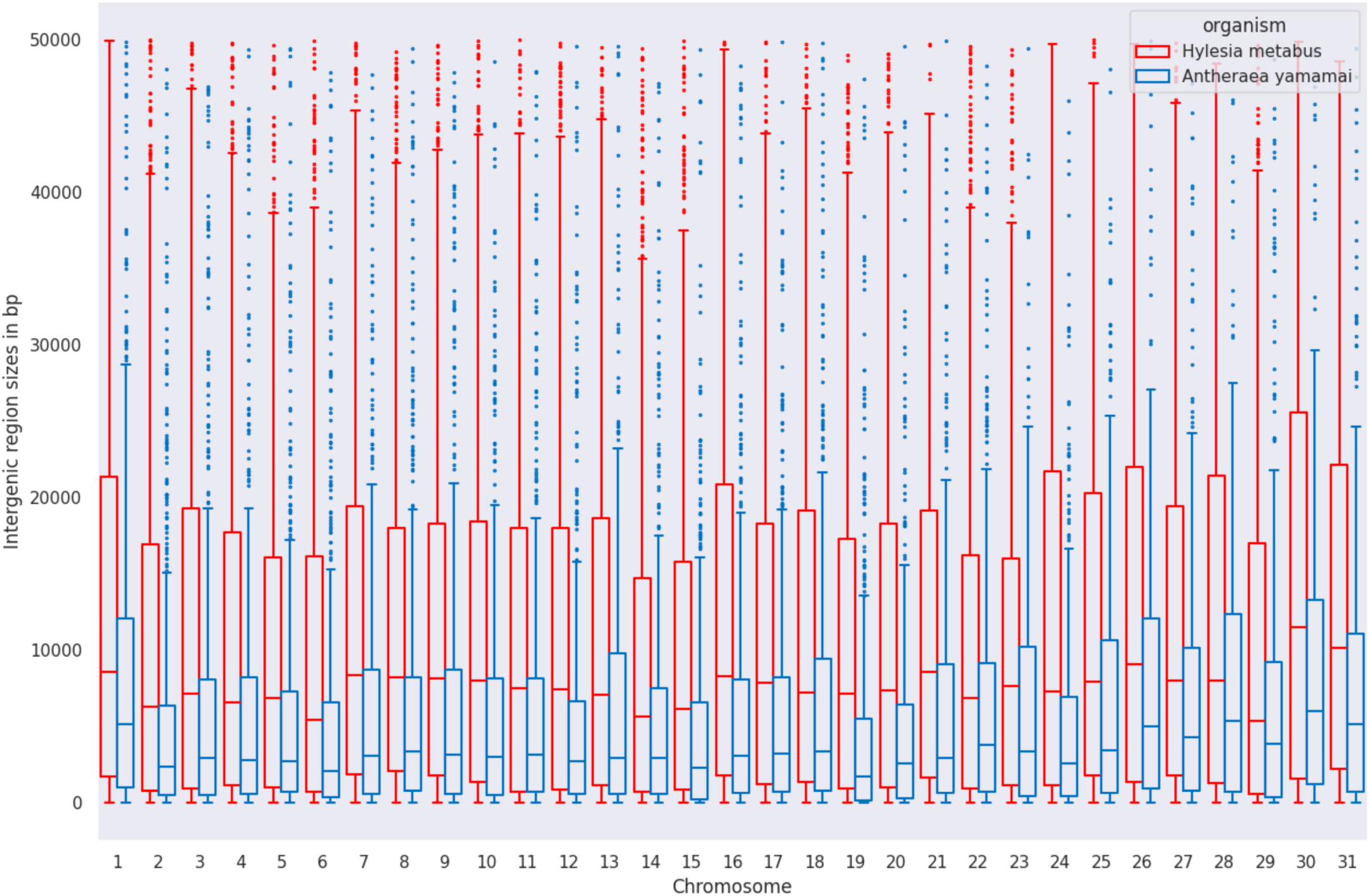
Distribution of Intergenic region sizes (measured in base pairs) for *Hylesia metabus* and *Antheraea yamamai* for the 31 chromosomes.

**Supplementary material 7.**
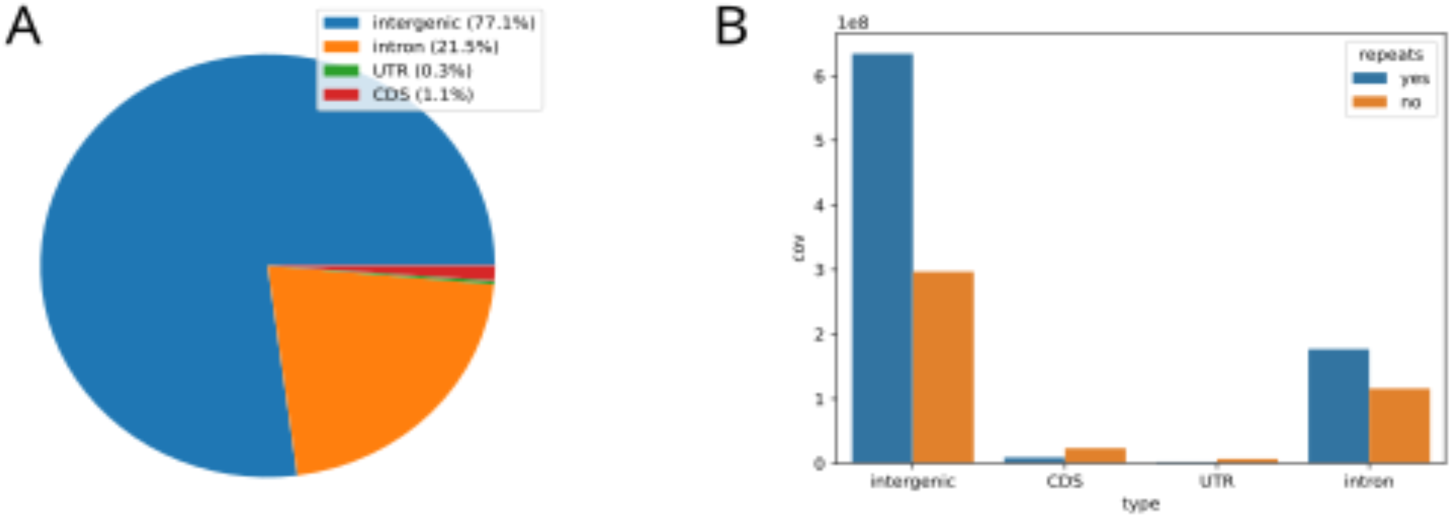
Location of repetitive elements in the *Hylesia metabus* genome **A.** Percentage of the coverage (measured in base pairs) of repetitive elements within the features (intergenic, introns, UTR, CDS) of the *H. metabus* genome. **B.** Coverage (measured in base pairs) of repetitive elements in genome features.

**Supplementary material 8.**
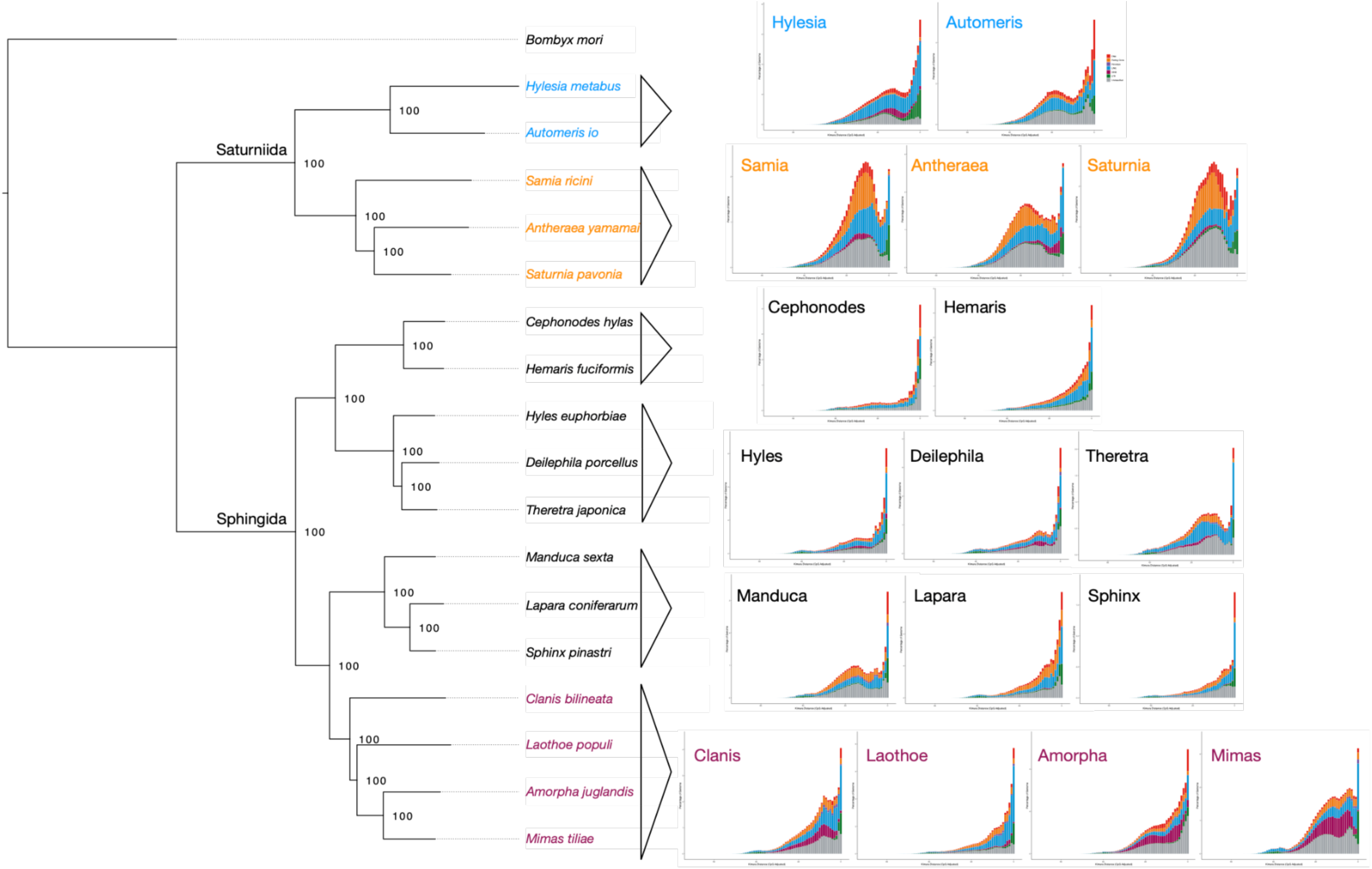
Comparison of the repeat landscape of *Hylesia metabus* versus the 17 other species considered.

**Supplementary material 9.**
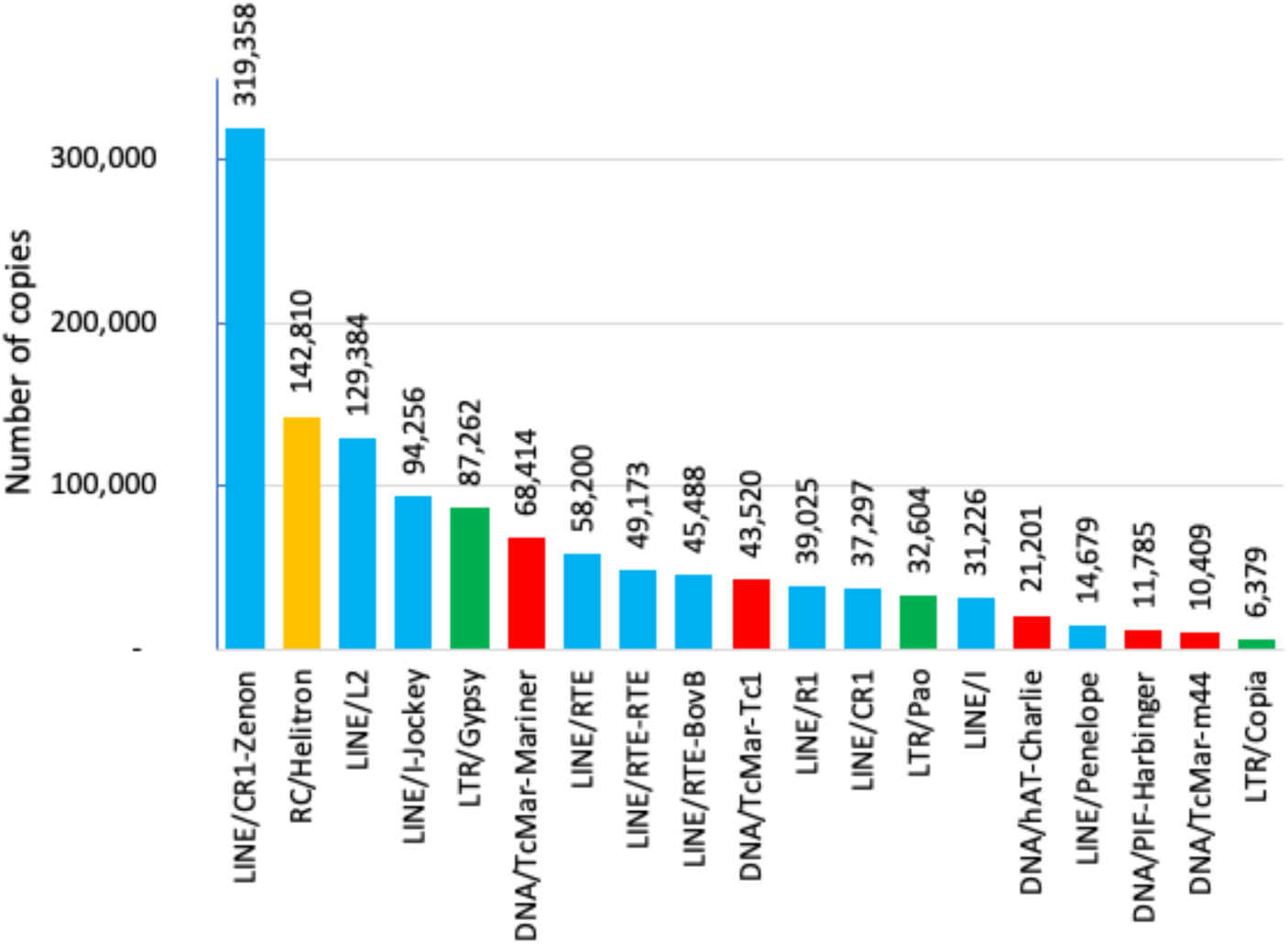
Number of copies for each of the most abundant TE families. Families are ranked by decreasing number of copies.

**Supplementary material 10.** Top TE families’ cumulative lengths in *Hylesia metabus* and corresponding cumulative lengths in the 4 other saturniidae species considered in the comparative analysis.

| TE family | *Hylesia metabus* | *Automeris io* | *Samia ricini* | *Antheraea yamamai* | *Saturnia pavonia* |
| --- | --- | --- | --- | --- | --- |
| LINE/CR1-Zenon | 109352923 | 20373335 | 2400252 | 17146553 | 7522389 |
| LTR/Gypsy | 82851380 | 8534372 | 3575502 | 4406707 | 4978918 |
| LINE/I-Jockey | 64188847 | 5329376 | 6890562 | 19504365 | 15290038 |
| LINE/L2 | 57568201 | 4639705 | 11209924 | 8357826 | 8187598 |
| LINE/R1 | 52633245 | 26144085 | 10099021 | 9859267 | 5723865 |
| RC/Helitron | 41902450 | 22080586 | 40561815 | 44592869 | 49941914 |
| DNA/TcMar-Mariner | 32879542 | 14149326 | 5783813 | 2101894 | 11188797 |
| LTR/Pao | 31997709 | 7264137 | 4308946 | 5949925 | 5995263 |
| LINE/I | 25512770 | 4603689 | 5006902 | 2463473 | 1816868 |
| DNA/TcMar-Tc1 | 24448122 | 4981093 | 1685254 | 3354857 | 3339704 |
| LINE/CR1 | 21468046 | 4794351 | 4224974 | 2889092 | 3136479 |
| LINE/RTE-RTE | 19156967 | 1622974 | 4310348 | 755187 | 2024669 |
| LINE/RTE-BovB | 17234452 | 2924644 | 7636678 | 1103102 | 6031714 |
| LINE/RTE | 15258135 | 5131841 | 4059525 | 260466 | 2659742 |
| DNA/TcMar-m44 | 9814187 | 167180 | 763528 | 64786 | 6982156 |

**Supplementary material 11.**
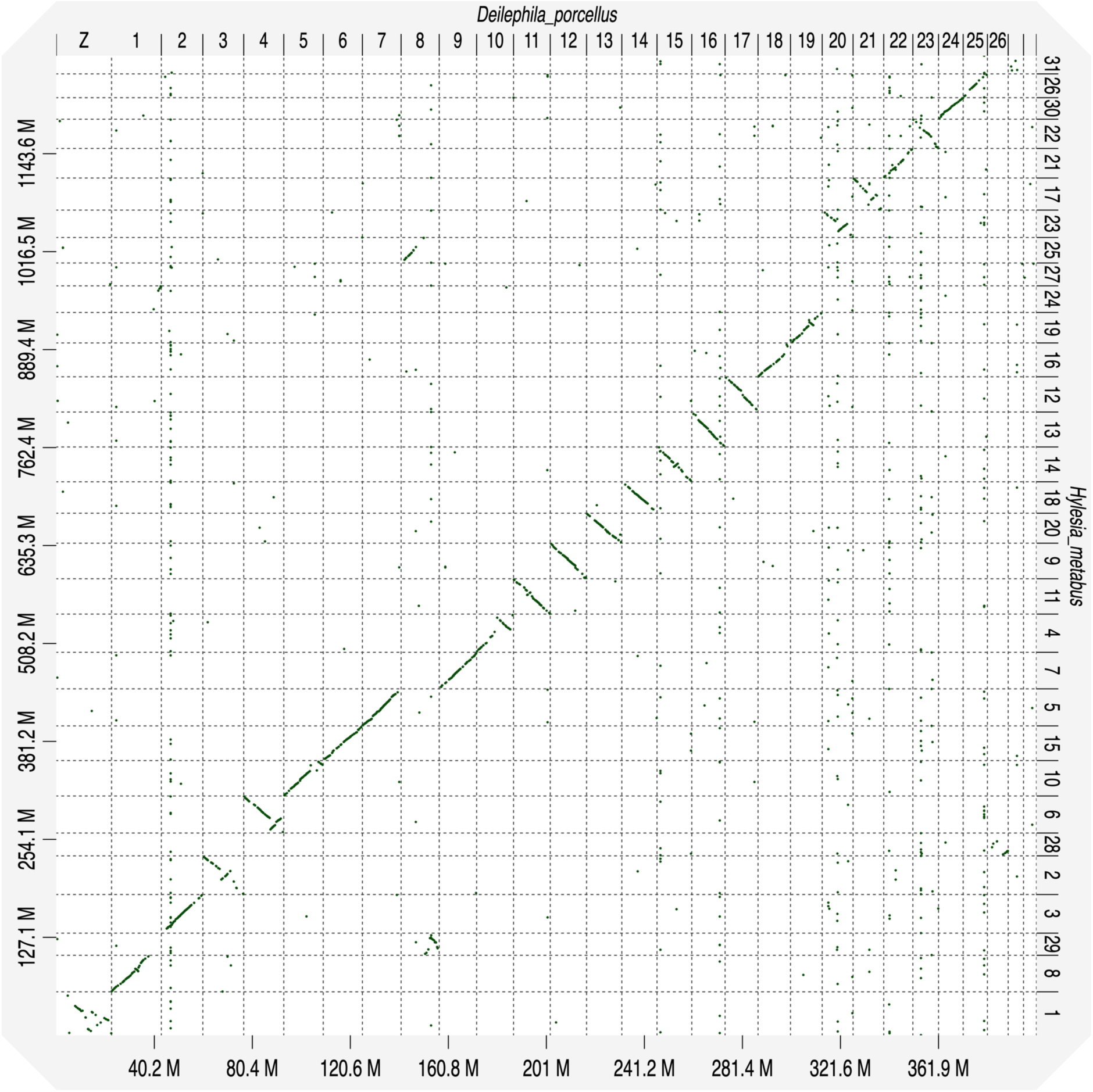
Synteny between *Hylesia metabus* and *Deilephila porcellus*. This graph notably illustrates that scaffold 1 in *H. metabus* corresponds to scaffold Z in *Deilephilia porcellus*.

**Supplementary material 12.**
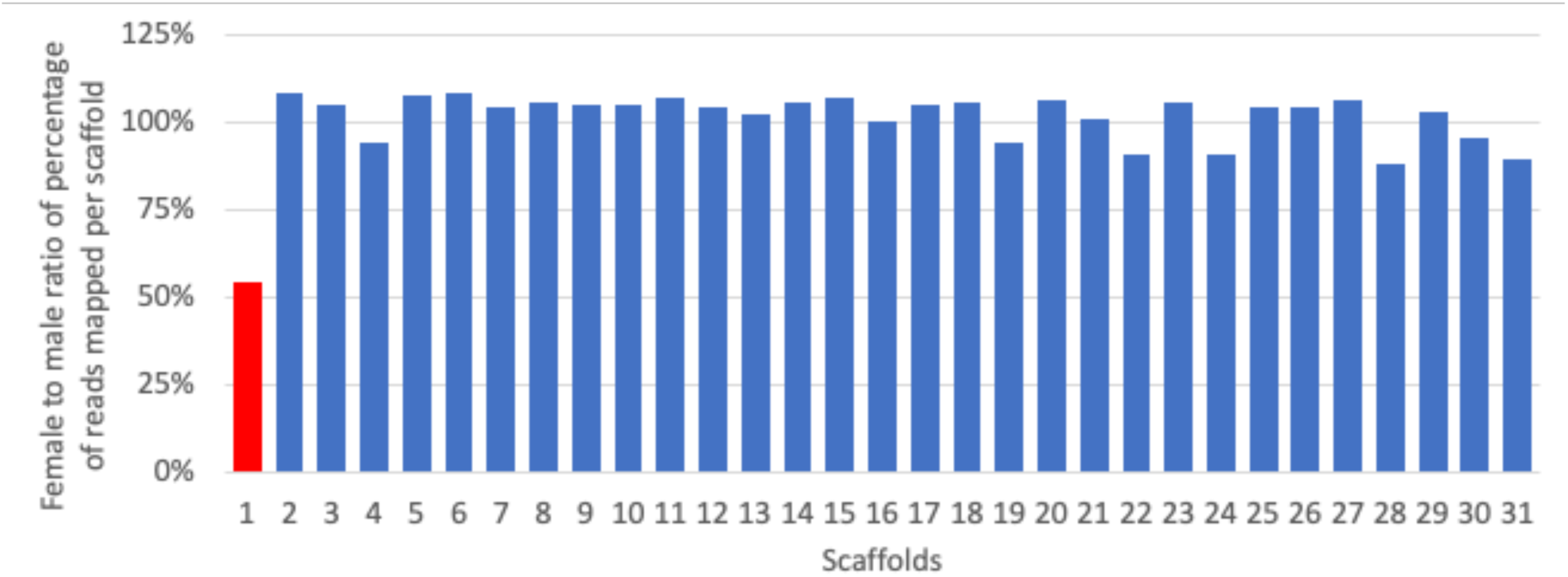
Female to male ratio of percentage of reads mapped per scaffold to identify the scaffold corresponding to the Z chromosome. The ratio of 54% on the scaffold 1 indicates that it has a single copy in females and hence corresponds to the Z scaffold.

